# Using relative brain size to better understand trophic interactions and phenotypic plasticity of invasive lionfish (*Pterois volitans*)

**DOI:** 10.1101/2020.01.10.902049

**Authors:** McKenna Becker

**Affiliations:** CIEE

**Keywords:** Cognition, crepuscular hunting, predator-prey interaction, relative brain mass

## Abstract

Predator-prey dynamics provide critical insight into overall coral reef health. It has been shown that predator-prey relationships link the relative brain size of predators to their prey. Predation pressure forces prey to use decision-making skills that require higher cognition by inspecting and identifying predators and then adjusting their behavior to achieve the highest chance for survival. However, the predation pressure that prey face outweighs the pressure predators face to find prey, resulting in prey having larger relative brain sizes than their predators. There is little data on the relative brain size of fishes with few natural predators such as *Pterois volitans*. This study compared the brain mass to body mass ratio of *Pterois volitans*, which have very few natural predators and thus very little predation pressure, to the brain mass to body mass ratio of their prey, possible predators, competitors, and taxonomically similar fish. Lionfish had a significantly smaller relative brain size than their predators, prey, and competitors, but was not significantly smaller than taxonomically similar fish. These results demonstrate that the morphological anti-predator adaptation of venomous spines causes little predation pressure. Thus, lionfish do not use the same cognitive skills as other prey or predators and, in turn, have smaller relative brain sizes.

## Introduction

Predator-prey relationships impact fish diversity, abundance, and distribution and can provide critical insight into greater coral reef health (Hixon and Beets 1993). Previous research has shown that an increase of predators introduced into a coral-reef can detrimentally affect the species richness and evenness of prey populations (Hixon 1986).

Interest has grown in examining predator-prey relationships through cognition and brain morphology. Kondoh (2010) furthered the understanding of predator-prey relationships by linking the brain mass to body mass ratio of predators to their prey. Kondoh (2010) analyzed 623 predator-prey relationships and found that 1) there is a strong correlation between log-scaled brain size to body size ratios of predators and prey, 2) predator-prey relationships are better identified when based on brain size to body size ratios, and 3) prey have a larger brain size to body size ratio than their predators. The question of how brain size to body size ratios can determine predator-prey relationships is better understood when taking cognitive ability and adaptive behavior (phenotypic plasticity) into account.

Predation pressure and other environmental changes induce learning and phenotypic plasticity within a generation of prey, which in turn maximizes their fitness (Kondoh 2010; Murren et al. 2015). Prey must be able first to inspect a fish and then identify it as a potential predator (Lima and Dill 1989; Murphy and Pitcher 1997). Furthermore, not every scenario with a predator is equally dangerous for prey, which means that prey must weigh the cost of energy to escape with the assessment of the risk of predation (Lima and Dill 1989; Ydenberg and Dill 1986). Once the prey has identified the fish as a possible predator and decided that the predator is a legitimate threat, it must adjust its behavior to achieve the best chance for survival (Murphy and Pitcher 1997). If predators must improve their ability to predate just as much as the prey have to improve their ability to escape, why do prey still have larger relative brains? Prey are under stronger pressure than predators because of what is called the life-dinner principle (Dawkins and Krebs 1979). The life-dinner principle states that with every predator-prey interaction, the prey will die if it makes the wrong decision. However, a predator will only need to find another prey (Dawkins and Krebs 1979). The risk assessments and decision-making processes due to greater predation pressure indicate that prey have higher cognitive ability and thus larger brain size to body size ratios than their predators (Kondoh 2010). However, in many cases, even predators were prey as juveniles, and can have predation pressure even into adulthood. As a result, predators can still have the same pressure to adapt their behavior to learn to avoid other predators. Kondoh (2010) hypothesized that as a prey’s anti-predator behavior improves, the predator must also improve its predation ability which results in a positive correlation between the brain size to body size ratio of predator and prey

*Pterois volitans*, commonly known as the red lionfish, has very few known natural predators in its native range of the Indo-Pacific as well as in its invaded range of the Western Atlantic (Albins and Hixon 2008). Lionfish have venomous dorsal, pelvic, and anal spines, which is likely why they have few predators (Allen and Eschmeyer 1973). However, whether they have a large or small relative brain size compared to their prey, predators, and other taxonomically similar species is still unknown. Since their invasion in the Western Atlantic, Gulf of Mexico, and Caribbean Sea, lionfish have been affecting the biodiversity, recruitment, and abundance of coral reef fishes (Albins and Hixon 2008; Green et al. 2012; Côté et al. 2013). Lionfish sightings were first reported in Bonaire, Dutch Caribbean in 2009, and as of 2015 are found at a higher density than in their native range (Green and Côté 2008; de Leon et al. 2013). Due to their lack of natural predators, divers began hunting lionfish almost immediately after they were first sited in Bonaire to mitigate the negative effects of this invasive species (de Leon et al. 2013).

A study by Côté et al. (2014) found that lionfish reacted sooner to divers after living in an environment where divers periodically culled for lionfish compared to lionfish that were not exposed to hunting divers. Claydon and Calosso (2012) found that not only were juvenile lionfish easier to capture by hand net than larger lionfish but also that adult lionfish actually swam away from divers to evade capture. The ability to recognize potential predators and act accordingly may present the possibility that lionfish have cognition and brain mass to body mass ratio similar to other prey fish. However, there should be a distinction between the predation pressure on a lionfish from a diver with a spear and the predation pressure of fish who have to learn to evade predators every day for survival. As such, although lionfish may learn to evade divers with spears, it may not present the same cognitive ability (and brain mass to body mass ratio) as other prey fish. Still, it could indicate a larger brain mass to body mass ratio than their predators.

This study investigated the brain mass to body mass ratio of invasive lionfish. Brain extractions of lionfish caught in Bonaire were conducted and compared to the brain mass to body mass ratio of their prey. This study also compared the brain mass to body mass ratio of lionfish to possible predators, competitors, and taxonomically similar fish which gave further insight to the predator-prey relationships of lionfish and the level of predation pressure that they face. To investigate the relationship between age (determined by length) and the ability to recognize divers as predators, this study analyzed the response of lionfish to divers with nets. There is little data about relative brain mass of fish that have few natural predators (i.e., lionfish) and how it relates to their cognitive ability. The results of this study give greater insight into using relative brain mass to better understand learning and adaptive behavior (phenotypic). The hypotheses of this study are the following:

H_1_: Lionfish will have a smaller brain mass to body mass ratio than their competitors because although they have similar prey, their competitors experience more predation pressure.

H_2_: Lionfish will have a smaller brain mass to body mass ratio than their prey

H_3_: Lionfish will have a larger brain mass to body mass ratio than their predators

H_4:_ Lionfish will evade divers when approached with nets

## Materials and methods

### Study site

Lionfish dissections took place at the CIEE Research Station in Kralendijk, Bonaire. Field data were collected on SCUBA dives at Yellow Submarine dive site. Lionfish used for lab data came from dive sites on Bonaire: Yellow Submarine, Bari Reef, Aquarius, Something Special, and Te Amo (Table 1). Yellow Submarine (12°9’36.20”N, 68°16’55.25”W) is located on the west side of Bonaire (Fig. 1). Yellow Submarine was chosen after preliminary dives showed the presence of lionfish at multiple depths. Due to the abundance of massive corals such as *Porites astreoides, Orbicella faveolata*, and *Orbicella annularis*, the structural complexity is favorable for lionfish because they are often found hiding in crevasses or under large overhangs when they are not hunting. Yellow Submarine is a fringe coral reef that is a popular site among divers to shoot lionfish which presents the potential for lionfish to learn to avoid divers. Yellow Submarine also has a lot of potential prey items for lionfish such as species of chromis, damselfish, wrasse, butterflyfish, and gobies.

**Fig 1.**
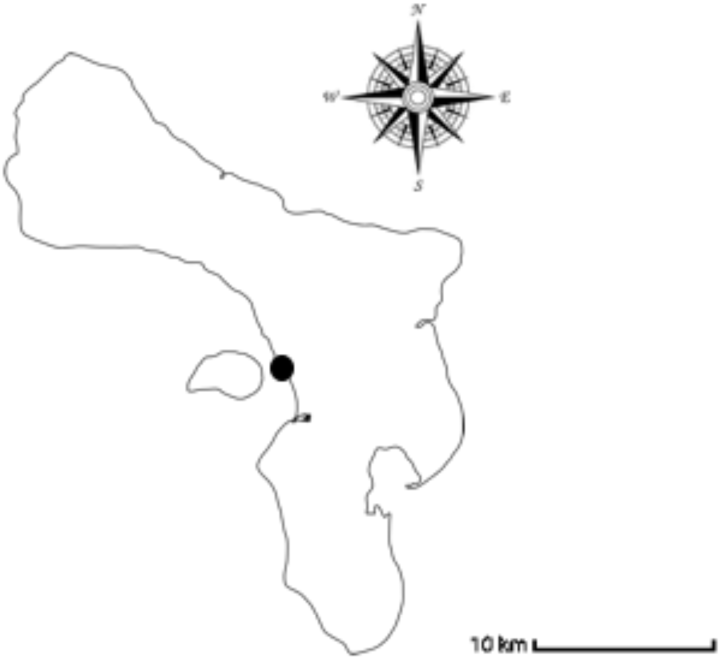
Map of Bonaire with the black dot indicating the Yellow Submarine dive site (12°9’36.20” N, 68°16’55.25” W) where field research was conducted.

### Study Organism

*Pterois volitans*, commonly known as the red lionfish, is native to the Indian Ocean and Western Pacific but has been found in the Western Atlantic and Caribbean Sea as an invasive species since 1985 (Schultz 1986; Semmens et al. 2004). Lionfish are found on the reef typically between 0 to 50m (Schultz 1985). Lionfish are crepuscular predators and are considered generalist consumers in both their native and invasive range (Morris and Akins 2009; Cure et al. 2012). With the exception of short periods during courtship, lionfish are solitary reef dwellers in their natural habitat and are usually found in crevasses or under overhangs (Schultz 1986). Lionfish are in the family Scorpaenidae (scorpionfish) which is characterized by having venomous spines (Fishelson 1975). Lionfish have 13 venomous dorsal spines, two venomous pelvic fins, and three venomous anal fins (Morris 2009). There are very few natural predators of lionfish that we know of, but there have been few instances where lionfish are found in the stomach content of larger groupers (Maljkovic et al. 2008).

### Lab methods

Lionfish used for dissection were shot and collected at various sites and depths in Bonaire between the years of 2013 and 2015 (Supplementary Table 1). Lionfish, as well as their brains, were extracted and weighed using a protocol created by the author (Supplementary Table 2).

### Field methods

Over the course of five weeks, six dives took place. Three of these dives occurred between 9:30-10:30 and three of these dives occurred between 18:00-19:00. These times were chosen because lionfish are crepuscular hunters meaning that they hunt during twilight hours in the morning and evening. Although the morning dives were after twilight hours, studies show that lionfish are still active in the morning after twilight (Côté and Maljkovic 2010). During each dive, I looked for lionfish for 30 minutes between 1-12 meters, and 30 minutes between 12-18 meters. During each dive I recorded 1) the length of each lionfish that I saw (1-10 cm, 11-20 cm, 21-30cm, etc.) 2) the depth of each siting and 3) their reaction to a diver swimming at them with a net. Behavioral responses were recorded as one of four categories; 1) swam away, 2) did not swim away, 3) defensive towards net, and 4) was hiding. Defensive behavior was recorded when the lionfish flexed their dorsal spines and intentionally swam toward the net or diver. To reduce the chance of observing the same individual on multiple dives, we conducted three dives headed north, each dive starting where the previous dive ended, and conducted three dives headed south using the same protocol.

### Data analysis

#### Lab data analysis

Lab dissection data specifically for lionfish were analyzed in two ways. First, every lionfish was plotted on a graph of ln[body size (g)] on the x-axis and ln[brain size (mg)] on the y-axis. This graph presents the overall trend of the brain mass to body mass ratio for lionfish found on Bonaire. A least-squares linear regression of brain size against body size was used to analyze correlation. Second, each lionfish was designated as either a juvenile (< 17.5 cm TL) or a sexually mature lionfish (≥ 17.5 cm TL) (Morris 2009) and plotted on a graph with body size (g) on the x-axis and brain size (mg) on the y-axis. A regression line was fit for each data set (juveniles and sexually mature lionfish). An analysis of covariance (ANCOVA) was used to determine whether the slopes of the two lines were significantly different. This analysis gave insight into whether juvenile lionfish had larger or smaller relative brain sizes than sexually mature adults.

The brain mass and body mass data of predators, prey, competitors, and taxonomically similar fish came from a study published by Michio Kondoh (2010). Kondoh (2010) published data for over 623 reef fish predator-prey pairs. Over 40 of those species from the families Pomacentridae, Apogonidae Acanthuridae, Serranidae, Scorpaenidae, and Labridae were used to analyze lionfish trophic interactions. Prey of lionfish were determined based on a study by Morris and Akins (2009) that investigated the stomach content of lionfish. The brain mass and body mass data of lionfish came from 26 dissections specifically for this experiment. To analyze relative brain size, every fish (lionfish, predators, prey, competitors, and taxonomically similar fish) was plotted on a graph with ln[body size (g)] on the x-axis, and ln[brain size (mg)] on the y-axis. A log-log least-squares regression of brain mass against body mass was used in order to give an equation that represented what the “predicted brain size” would be. The equation for predicted brain size was ln[brain size (mg)] = 0.6118 (ln[body size (g)]) + 2.7641 (Fig. 3). Finding the predicted brain size allowed me to address the confounding factor that larger fish have larger brains whether they are a predator or prey. Once the predicted brain size for every species was found, it was subtracted from the actual brain size to find a residual. The residual represents how much the actual brain size deviates from the predicted brain size for a given body size fish. The more negative the residual, the smaller the relative brain; the more positive the residual, the larger the relative brain. The residual (referred to from this point forward as relative brain size) was used to compare the brain sizes between lionfish and their predators, prey, competitors, and other taxonomically similar fish.

**Fig. 2.**
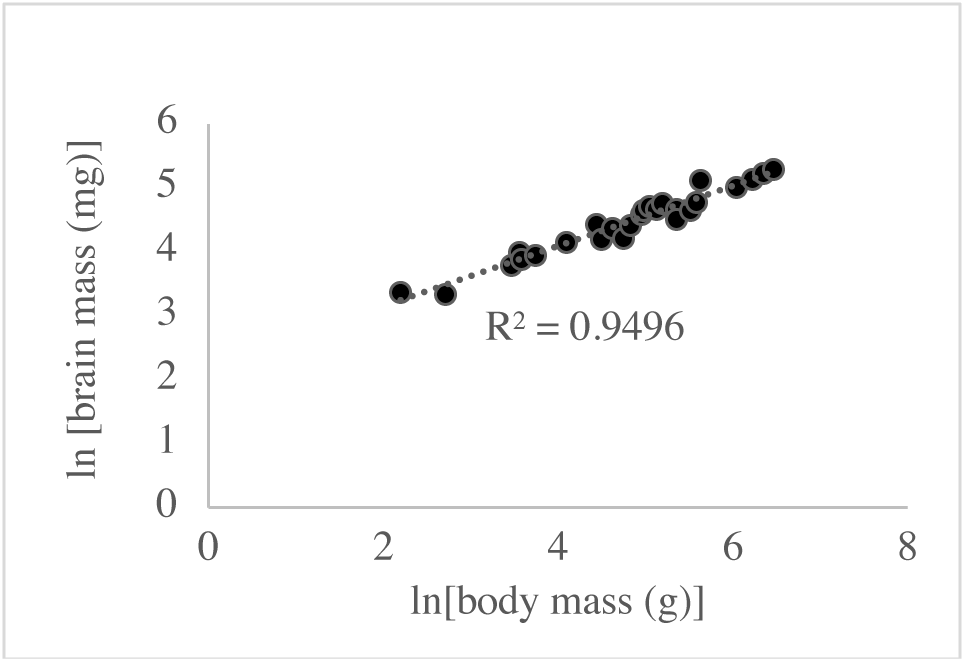
Log scaled brain mass to body mass ratio of lionfish revealing that age did not affect relative brain size.

**Fig. 3.**
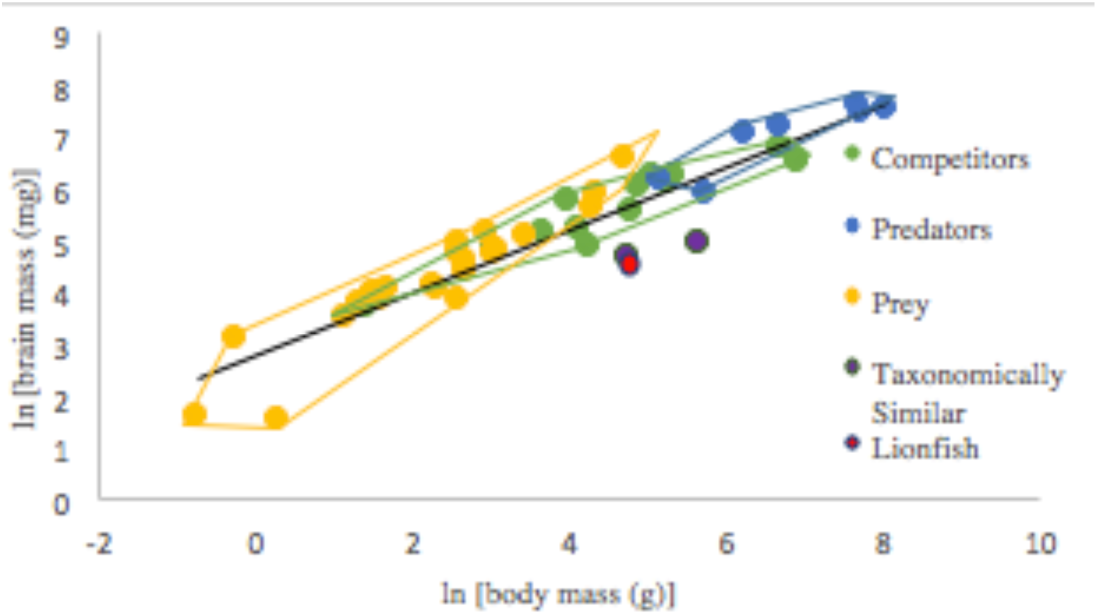
The relationship between ln[body mass (g)] and ln[brain mass (mg)] of lionfish to possible predators (n=7), prey (n=22), competitors (n=14), and taxonomically similar fish (n=2). Lionfish have the smallest relative brain size. Colored lines represent the boundaries of the most extreme data points.

#### Field Data Analysis

Behavioral response of lionfish to divers with nets was the primary focus of the field data collection. The response of lionfish was compared to their length with the assumption that length and age are related. This data was presented on a clustered bar graph. Total length (cm) represented each bin on the x-axis and the amount of times that the lionfish displayed each response to the diver on the y-axis. Within each bin, individual bars represented each behavior. This graph allowed me to compare total length with the frequency of each behavior. However, the data presented by this graph indicated that age did not have a significant impact on their behavioral response. Thus, the data was pooled together on a graph that presented all of the lionfish together on a bar graph with the behavioral responses on the x-axis and the percentage of the time that they displayed that behavior on the y-axis.

## Results

### Relative brain size of juvenile and sexually mature lionfish

There was a positive correlation between log-scaled brain mass and body mass of lionfish (R^2^ =0.94964) (Fig. 2). The brain mass to body mass ratio of sexually mature lionfish (>17.5 cm TL) (Morris 2009) was not significantly larger than the brain mass to body mass ratio of juvenile lionfish (<17.5 cm TL) (p = 0.125, d.f. = 1). Overall, 26 lionfish were dissected including 7 juveniles and 19 sexually mature lionfish.

### Lionfish have smaller relative brains than their prey, predators, and competitors

Lionfish had the smallest relative brain size when compared to the averages of all other functional groups (Fig. 3 and 4). Lionfish have a significantly smaller relative brain size (−1.248) than their prey (0.1044 ± 3.649-) (mean ± standard deviation) (p < 0.0001, d.f. = 1, 21, t = 12.94) and their predators (0.0804 ± 0.2902) (p < 0.0001, d.f. = 1, 6, t = 12.12) (Fig. 3 and 4). Lionfish also have a significantly smaller brain size than their competitors (0.0487 ± 0.3059) (p < 0.0001, d.f. = 1, 13, t = 15.86) (Fig. 3 and 4). There was no significant difference between the relative brain mass of lionfish and taxonomically similar fishes (−1.449 ± 0.2308) (p = 0.640, d.f. = 1, 1, t = 0.63) (Fig. 3 and 4).

**Fig. 4.**
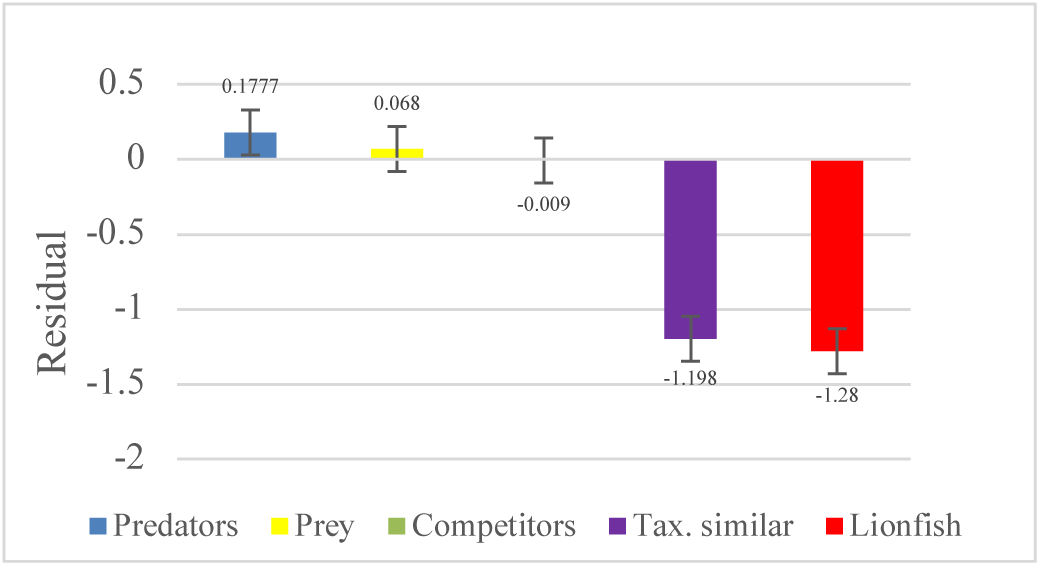
The residual=(ln(actual brain size)) - (predicted brain size) for predators (n=7), prey (n=22), competitors (n=14), and taxonomically similar fish (n=2). Negative residuals indicate their average brain size is smaller than their predicted brain size.

### Behavioral response to divers with nets

Lionfish displayed similar responses to divers regardless of their total length (TL). Lionfish that were 1-10 cm TL were found to swim away from divers 50% of the time and not swim away from divers 50% of the time (n = 4). Lionfish that were 11-20 cm TL swam away from divers 60% of the time, did not swim away from divers 13% of the time, were defensive towards divers 13% of the time, and were found hiding 13% of the time (n = 15). Lionfish that were 21-30 cm swam away from divers 50% of the time, did not swim away from divers 25% of the time, and were aggressive towards divers 25% of the time. Length of lionfish did not have a substantial effect on behavior towards divers. Thus, I decided to pool all of the lionfish together to analyze the data (Fig. 5). Lionfish swam away from divers 56% of the time, did not swim away from divers 21% of the time, were aggressive towards divers 13% of the time, and were hiding 8% of the time (n = 23).

**Fig. 5.**
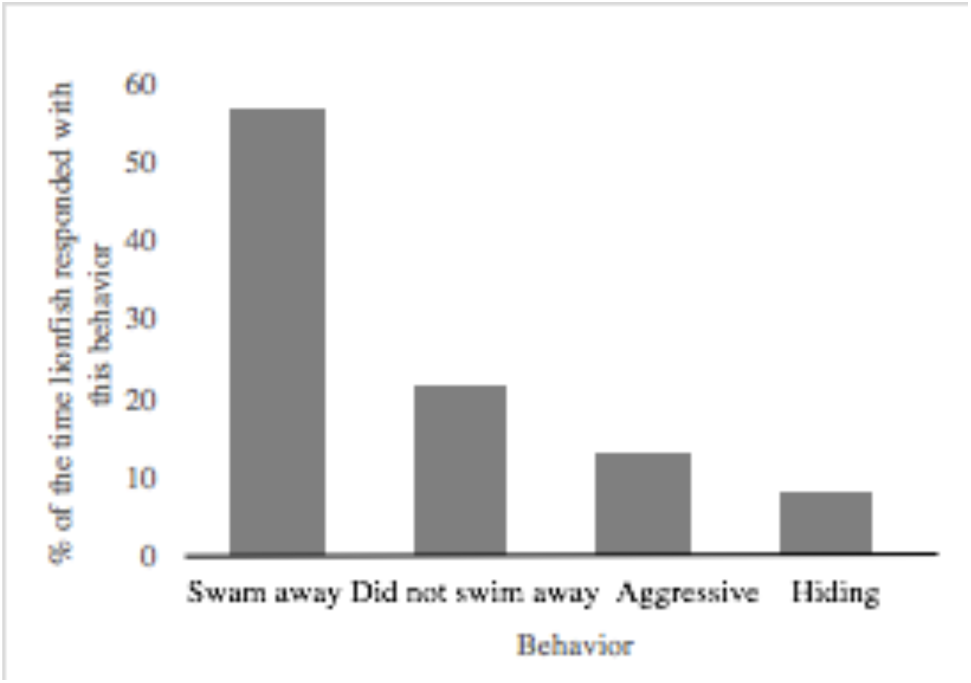
The comparison of different behavioral responses by lionfish to divers swimming at them with nets over the course of 6 dives in 5 weeks. Lionfish swam away from divers 57% of the time, did not swim away from divers 22% of the time, were aggressive towards divers 13% of the time, and were hiding 8% of the time (n = 23).

## Discussion

Lionfish had significantly smaller relative brain size than their predators, prey, and competitors. However, they did not have a significantly smaller brain size than taxonomically similar fish. These results support H_1_ and H_2_ which stated that lionfish would have smaller relative brains than their competitors and prey. The results of this study did not support H_3_ which stated that lionfish would have larger relative brains than their predators. Lionfish did swim away from divers 56.5% of the time which, indicates that lionfish may have adapted to recognize divers as predators, but the predation pressure is not great enough to change their gross morphology. However, their small brains are mostly likely due to evolutionary constraints which is evident due to other scorpaenids that also have small brains.

Lionfish and other Scorpaenidae fish (scorpionfish family) had very similar relative brain sizes, which suggests that it is a taxonomic constraint rather than predator-prey interactions that cause lionfish to have relatively small brains. Predation pressure is a large component in forming morphological anti-predator adaptations, such as venomous spines (Lima and Dill 1989). Lionfish and other scorpionfish that were used in this study have venomous spines which are used as defense in result of an anti-predator morphological adaptation (Halstead and Chitwood 1955; Casewell et al. 2013). Thus, lionfish do not have to use cognition to maximize fitness because they rely on their morphology.

Lionfish morphology can also explain why they have smaller relative brain sizes than their predators. Even some predators are prey at some point when they are juveniles and thus must still use decision making processes to maximize their fitness. However, due to lionfish’s venomous spines, even as juveniles they do not face the same predation pressure that other juvenile predators do. The lack of predation pressure due to their venomous spines at all life stages never allow lionfish to learn how to identify and avoid natural predators using the same nuance that other prey have to use. However, lionfish did avoid divers with nets as predators 56% of the time (70% of the time when aggression is included as an anti-predator behavior) which indicates that they are capable of some phenotypic plasticity. The probability of being killed during a period of time is called the risk of predation (Lima and Dill 1989). Thus, the low abundance and frequency of hunting divers may apply a smaller predation pressure onto lionfish (due to lower risk of predation) than other predators do onto other prey, and a smaller predation pressure than other predators feel when they are juveniles. Comparing the brain mass to body mass ratio of lionfish to their competitors who have similar prey but may experience more predation pressure gave insight into the degree that predation pressure effects brain mass.

A possible limitation of this study could be in comparing the brain mass of lionfish with every other fish due to different extraction processes. The brain mass and body mass data for fish from the Kondoh (2010) study came from FishBase but the method for extracting the brains was not stated.

Lionfish morphology may be a fundamental limitation to their phenotypic plasticity. Anti-predator mechanisms can be behavioral or morphological (Lima and Dill 1990) but the findings of this study suggest that only behavioral anti-predator mechanisms contribute to larger relative brain sizes than their predators. Further studies should include investigating how morphological anti-predator mechanisms, such as venom, affect their ability to adapt to introduced predators in a controlled environment.

## Acknowledgements

I would like to thank my research advisor Dr. Patrick Lyons and my co-advisor Nathaniel Hanna Halloway for their insight and support. I would also like to thank Dr. Rita Peachey and the rest of the CIEE staff for making my project possible.

## Supplemental Tables

**Supp. Table 1.** List of dive, their latitude and longitude, number of lionfish shot at which depth and what year. Six lionfish were shot at a dive site that was not recorded. The depth that lionfish were shot at Yellow Submarine and Something Special was also not recorded.

| Site name | Latitude, Longitude | # lionfish shot at site | Depth lionfish were shot (m) | Year |
| --- | --- | --- | --- | --- |
| Aquarius | 12°5'25.21"N,<br>68°16'59.84"W | 9 | 8-28 | 2015 |
| Bari Reef | 12°9'48.67"N,<br>68°17'13.59"W | 1 | 30 | 2015 |
| Something Special | 12°9'45.56"N, 68°17'6.86"W | 3 | Unknown | 2015 |
| Te Amo | 12°8'3.94"N, 68°16'47.66"W | 4 | 15-20 | 2013 |
| Yellow Submarine | 12°9'36.20"N,<br>68°16'55.25"W | 4 | Unknown | 2015 |
| Unknown |  | 6 |  |  |

**Supp. table 2.**
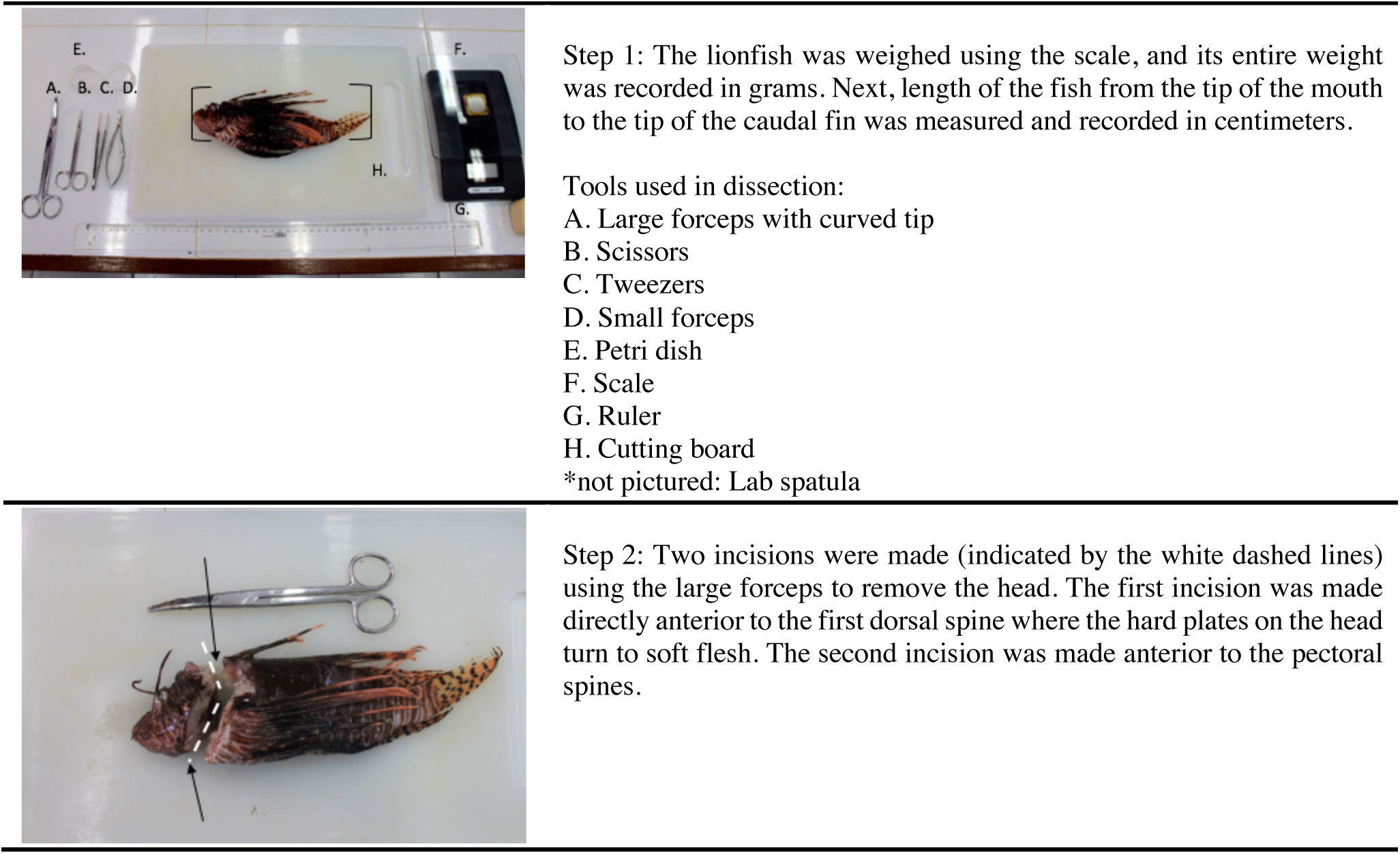
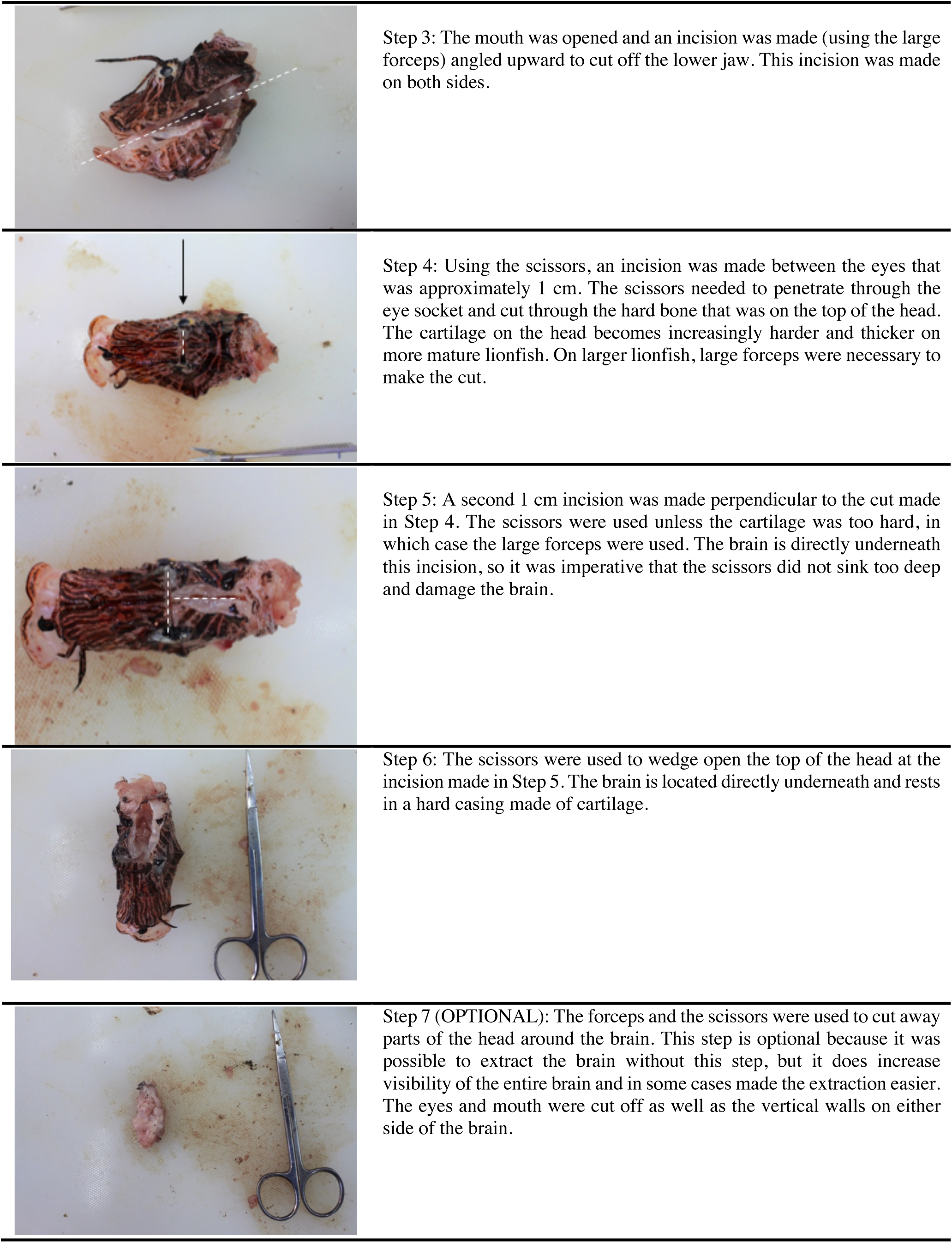
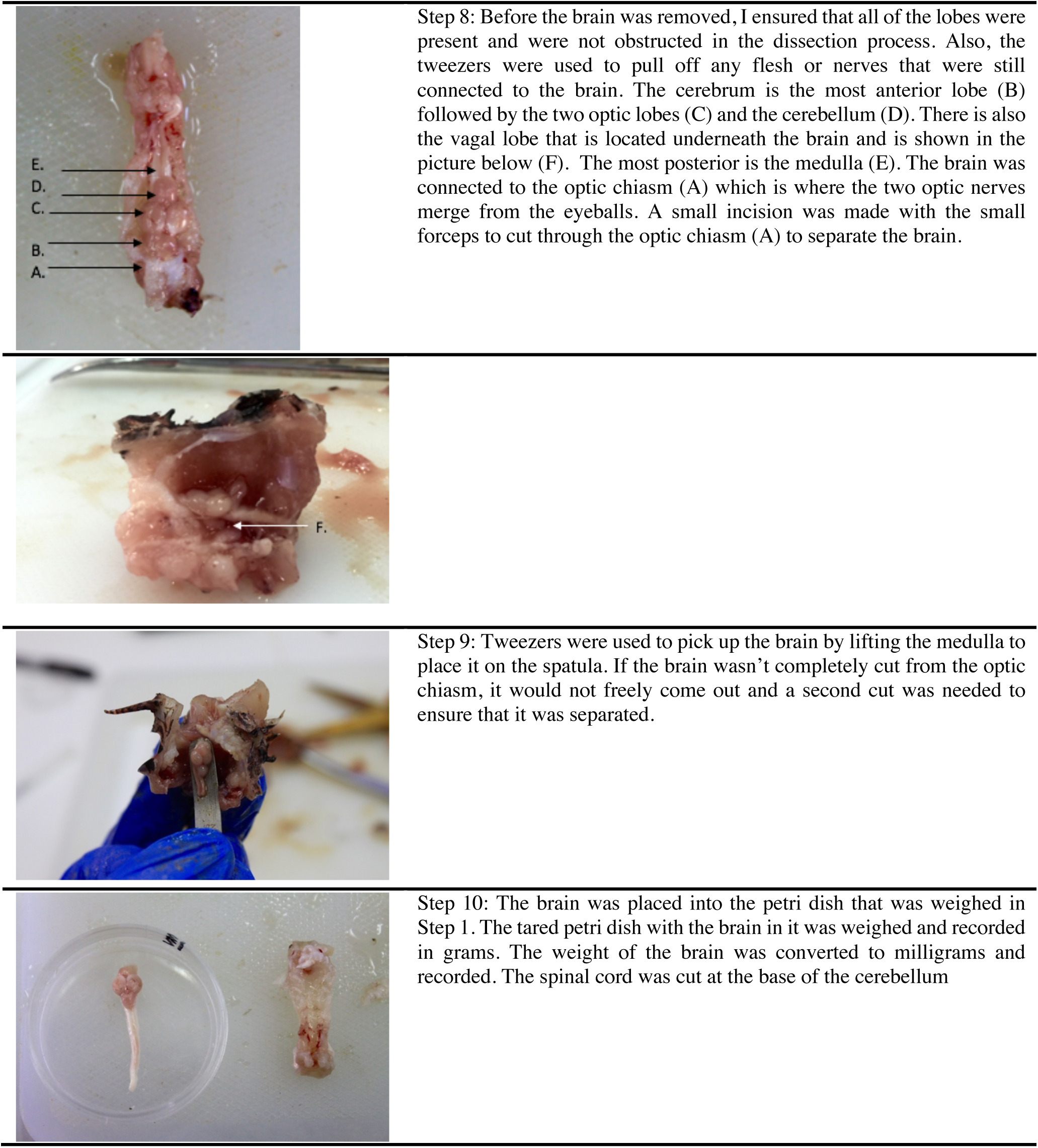
Step by step procedure for extracting a lionfish brain and the exact tools needed for dissection (Step 1) as well the neuroanatomy of a lionfish brain (Step 8). The protocol for this procedure was created by the author and specifically for the current study.

